# Environmental microbial extracts for longitudinal studies of gut microbiome assembly and maintenance

**DOI:** 10.64898/2026.06.12.732002

**Authors:** Rahul Bodkhe, Rebecca Choi, Michael Shapira

## Abstract

Animals harbor diverse gut microorganisms that influence host health and fitness. Synthetic microbial communities have been instrumental in enabling reductionist studies of host-microbiome interactions, but some questions require microbial communities with more natural-like complexity while preserving experimental tractability, *in vivo* monitoring, and quantitative analysis. Here, we describe a method optimized for longitudinal studies of host-microbiome-environment interactions in the nematode *Caenorhabditis elegans*. In this approach, complex microbial extracts (CMEs) are generated from environmental samples and applied to worm culture plates, providing a diverse yet experimentally convenient microbial environment. We show that CME composition remains stable during cold storage, enabling reproducible longitudinal experiments while minimizing confounding environmental drift over time. As a proof of principle, we apply this method to examine age-dependent changes in the worm gut microbiome, providing support for previous reports of age-dependent increase in the abundance of gut *Enterobacteriaceae*. CMEs provide a practical and reproducible framework that complements experiments using monocultures or synthetic communities, enabling longitudinal studies of host–microbiome interactions under conditions that better approximate natural microbial complexity.

## Introduction

It is now well appreciated that most if not all animals harbor complex gut bacterial communities [1]. Over the past two decades, research has highlighted the critical roles of these communities in animal health, evolution, and ecological interactions [2–6]. Simple model organisms such as *Drosophila melanogaster*, Zebrafish and *Caenorhabditis elegans* have emerged as powerful models for interrogating fundamental aspects of host-microbe interactions [7–11]. Their genetic tractability, short generation times, and well-characterized biology enable experimental manipulation and mechanistic analysis. In *C. elegans*, for example, microbiome research has advanced understanding of microbial contributions to infection resistance, aging, transgenerational inheritance, and behavior [12–16]

In nature, animals are continuously exposed to diverse microbial communities yet maintain characteristic gut microbiomes that represent only a subset of the surrounding microbial diversity, acquired through feeding, environmental contact or social interactions. *C. elegans*, for example, inhabits microbe-rich niches such as rotting fruit and decomposing plant material [5,17–20], where it encounters highly complex and dynamic microbial communities, yet maintain a gut microbiome distinct from its environment [21–23].

To model host-environment interactions under controlled conditions, microbiome studies in *C. elegans* employed defined synthetic communities spanning different levels of diversity, including CeMbio [22] SC20 [24], and BIGbiome [25], composed of bacterial gut isolates representative of the worm’s native microbiome. Similar strategies have been applied across a range of host systems, with host-specific synthetic communities (SynComs) developed for *Drosophila* [26], mosquitos [27], honeybees (Kešnerová et al. 2017), as well as for vertebrate systems [29]. These approaches offer experimental control and reproducibility but necessarily simplify the complexity of natural microbial environments. Conversely, more naturalistic systems such as environmental microcosms can better capture ecological complexity but but present practical challenges in sampling, observation, and quantification [21,30–33]. This trade-off is particularly pronounced in hosts with highly complex microbiomes, where simplified SynComs may inadequately represent the diversity and functional capacity of native microbial communities.

Here, we describe a complementary approach for microbiome research based on complex microbial extracts (CMEs), which preserve natural-like microbial diversity while enabling reproducible sampling, quantification, visualization, and long-term storage suitable for longitudinal studies. We demonstrate the utility of this method by applying it to characterize age-dependent changes in the gut microbiome of *C. elegans*. CMEs therefore provide an experimentally tractable intermediate between simplified synthetic communities and ecologically complex microcosm approaches for studying host–microbiota interactions.

## Methods

### Strains

Wildtype worms of the N2 strain were used in all the experiments. Worms were raised at 20°C on OP50 or CME on peptone-free medium (PFM) to minimize bacterial growth and competition once seeded on plates.

### Compost microbial extracts (CMEs)

Soil samples were collected from different locations across the Berkeley campus. To enrich and diversify bacterial communities, collected soils were supplemented with organic matter in the form of chopped apple, banana peels, or a combination of both and composted at room temperature for around two weeks, mixing every other day.

Preparation of microbial extracts was initiated by mixing 45-50 grams of compost with 20-30 mL M9 buffer in a 50 mL conical tube, followed by vigorous vortexing (Fig. 1A). Soil particles were then pelleted by a two-minute centrifugation at 1800 rpm. Lower-density particles (and native worms, if present in the original soil samples) were further removed by filtering the supernatant through a double layer of sterile gauze, and the resulting extract, typically around 20 mL, was collected in a separate conical tube. This procedure was repeated with more compost until ∼ 200 mL of extract were collected. Filtered extract was aliquoted into 1.5 mL microfuge tubes and centrifuged at 10,000 rpm for two minutes to pellet bacteria. Supernatant was discarded and CME pellets were combined to double the quantity per tube, air dried for 30 minutes and stored at 4°C until use. When used in experiments, CME pellets were resuspended in 200 µL M9, plated on PFM plates, and used as a source of food and commensals for approximately 100 worms, which were monitored to ensure development to maturity.

**Figure 1.**
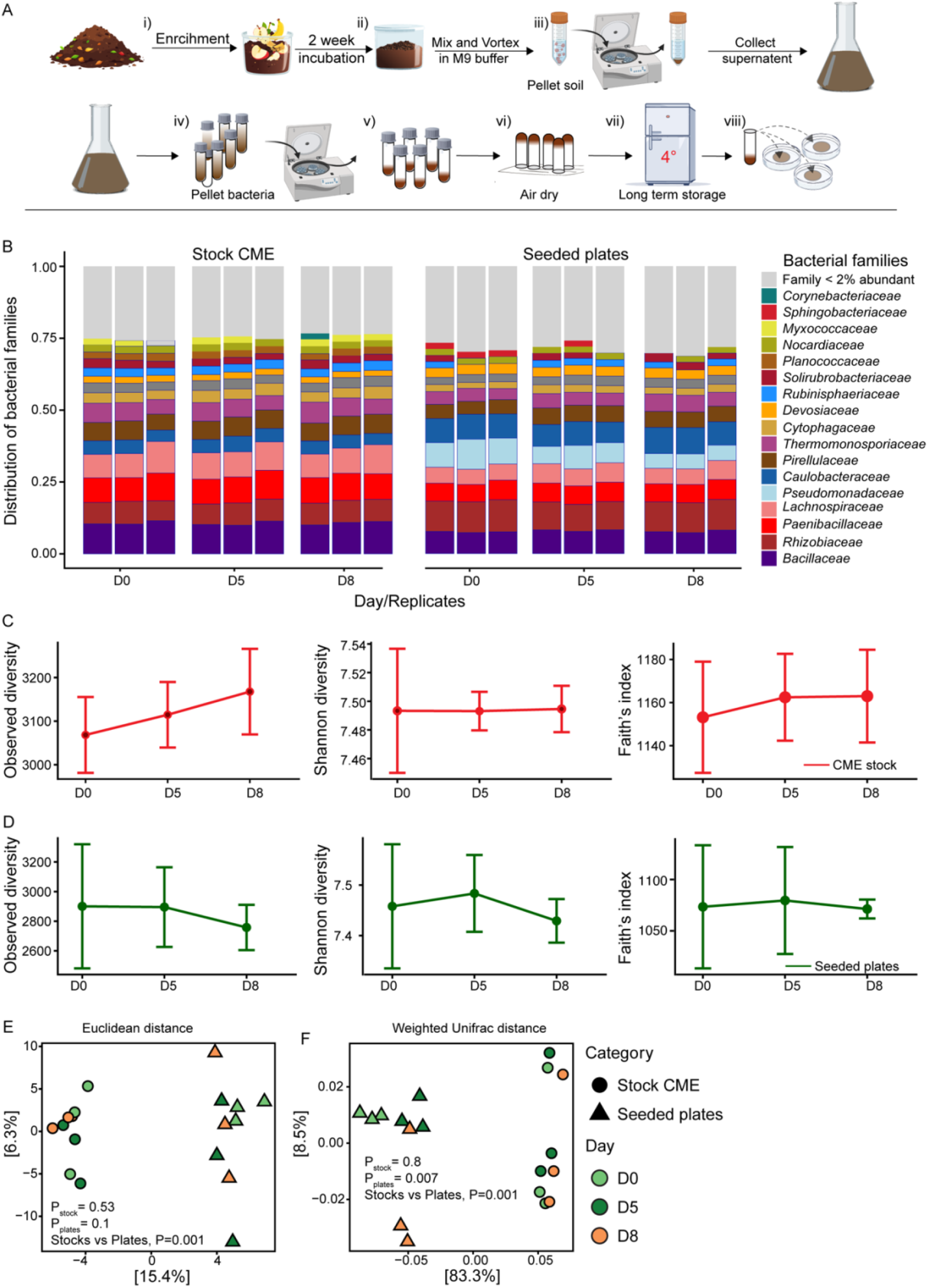
Compost microbial extracts maintain a consistent microbiome composition for microbiome research. **(A)** Schematic depiction of CME preparation. (i) Compost soil was enriched with produce to enhance microbial diversity. (ii) After two weeks of incubation, soil and dense particles were removed by filtration and low-speed centrifugation. (iii–v) Bacterial biomass was concentrated by high-speed centrifugation. (vi) The resulting pellet was air-dried and stored for long-term use. (vii) CME can be conveniently used by plating on PFM plates. **(B)** Relative abundance of bacterial families in CME stocks and on plates 24 hours after seeding. Each stacked bar represents an independent CME sample, color-coded by bacterial family. **(C)** Observed richness, Shannon diversity, and phylogenetic diversity of stored CME stocks analyzed at designated days **(D)** Similar alpha diversity indices for CME communities on seeded plates. Data are shown as mean ± SD from three independent samples (N = 3 per time point). Differences between time points were found to be non-significant (Welch’s t-test). **(E)** Principal component analysis based on Euclidean distance, or (**F)** weighted UniFrac distances, clustering microbiomes in CME stock samples or plates (p-values obtained with PERMANOVA).

### Worm aging experiment

Aging experiments were carried out essentially as in [24]. Briefly, synchronized worm populations were prepared by bleaching gravid worms and releasing eggs that hatched in M9 salt solution to provide germ-free L1 larvae. L1 worms were raised at 20^°^C on plates freshly seeded with CME, transferred daily to fresh plates to separate the original cohort from their progeny, enabling the continuation of experiments into what could be considered as middle age.

### Sampling for microbiome analysis

Samples of CME stock communities kept as aliquots at 4^°^C, were taken out of storage in designated time points and either used for DNA extraction or resuspended in M9 and seeded onto PFM plates from which they were harvested the following day. Samples were collected and kept at -20^°^C until DNA extraction.

Gut microbiomes of aging worms were analyzed in synchronized worm populations raised on CME-seeded plates. Worms were washed off with M9 into 15 ml conical tubes and pelleted by light centrifugation (1800 rpm) on a clinical centrifuge. Following aspiration of supernatant, worms were transferred onto the center of a sterile 100 mm 2% agar plate, allowing worms to disperse away from the any remainders of grazed CME. The central agar region containing grazed CME was excised and discarded. Worms from the remaining plate were washed off and collected in 15 mL falcon tube using M9 buffer + 0.0125% Triton X-100 (M9 + T). Worms further washed three times with M9 + T, paralyzed with 10 mM levamisole to prevent pharyngeal pumping, and surface-sterilized with 2% bleach in M9 [23]. Worms were washed twice with sterile M9 to remove bleach and stored at -20°C until DNA extraction.

### DNA extraction for 16S rRNA gene sequencing

DNA was extracted from worm and CME samples as previously described, using the DNeasy PowerSoil Pro Kit (Qiagen Cat. #47016) and eluted in 60 μl elution buffer [24,34]. DNA extracted from worm and CME samples were used as a template in PCR reactions with primers 515F and 806R and with Illumina Nextera XT DNA Library Preparation Kit (Cat. #FC-131-1096) to generate sequencing amplicon libraries of the bacterial 16S rRNA V4 region, as previously described [34]. Paired-end sequencing was performed on an in-house Illumina Miniseq platform using the High-Output (300 cycles) Illumina kit (Cat. #FC-420-1003).

### Microbiome analysis

High throughput Sequencing yielded high quality data (Q30) with an average of 109482 ± 29212 reads per sample (Supplementary Table 1). FASTQ files were processed using DADA2 v1.32 [35], including estimation of error models, denoising, merging of paired end reads and removal of chimeras (using the Consensus method) to generate a table of amplicon sequence variants (ASVs) containing an average of 61666 ± 21911 reads per sample. Subsequently, ASVs with fewer than 9 total reads across analyzed samples were excluded. Taxonomic assignments were carried out using the SILVA database genus (v138.1) [36].The final ASV table, merged with the corresponding metadata and taxonomy files were imported into R using phyloseq v1.50.0 [37] for downstream analyses. Raw data is available in the NCBI SRA database with accession number PRJNA1462967.

#### Alpha Diversity

Observed diversity and Shannon index were calculated using the plot_richness() function in phyloseq. Faith’s Phylogenetic Diversity (PD) was calculated using the Picante package.

#### Beta diversity

was evaluated using complementary distance metrics capturing abundance, compositional structure, and phylogenetic relationships. Euclidean distances were calculated following centered log-ratio (CLR) transformation, and weighted UniFrac distances were calculated using phylogenetic tree data using the distance() function in phyloseq. The phylogenetic tree was constructed by aligning sequences with DECIPHER [38] and building a neighbor-joining tree using phangorn [39]. Differences in community composition were tested using Permutational Multivariate Analysis of Variance (PERMANOVA) implemented in the adonis2() function of the vegan package with 999 permutations.

#### Taxonomic Composition

Stacked bar plots were generated using relative abundance data aggregated at the bacterial family level to visualize community composition across samples. Graphs were drawn using the ggplot2 R package [40]. R-scripts used to generate figures are available at https://rpubs.com/MicroRB/CME_Method_Paper

## Results

### CME compositional stability

To examine whether stored CME can serve as a stable microbial source for longitudinal microbiome studies, we analyzed the microbiome composition of a concentrated microbial extract (CME) prepared with soil composted with apples and banana peels that was stored at 4°C over a week (Fig 1A). CME aliquots, resuspended in M9, were used for DNA extraction and 16S analysis immediately from storage, or used to seed plates, and harvested from plates for DNA extraction one day later, examining community composition after settling on plates (Fig 1B). No apparent difference was observed in community composition of CME samples over a period of over a week (Fig. 1B left). Analysis of community composition on seeded plates may expose more nuanced changes, as growth on plates (which is minimal on the peptone-free media, but not completely negligible) may allow inter-bacterial competition resulting in small changes in the original stock that are then amplified by competition. Indeed, communities on seeded plates have somewhat changed during the 24 hours, compared to the original stock, which primarily involved expansion of *Pseudomonadaceae* bacteria, similar to previous reports [6]. Yet, community composition in seeded plates largely remained stable over the eight-day course of CME sampling.

In analysis of microbial diversity performed on CME stock samples shown in Fig. 1B, all alpha diversity indices showed no significant change during the eight-day storage (Welch t-test, p > 0.05 Fig. 1C). A similar trend was observed in seeded plates (Fig 1D). Principle component analyses, assessing beta diversity, highlighted the differences between CME stock samples and their plate-settled version (Fig. 1E and F), but again showed no significant changes in community composition in longitudinally sampled CME stock aliquots, assessed either with Euclidean distance or with weighted UniFrac distances. While some statistically significant differences were found among seeded plates communities, this was apparent only with weighted UniFrac distance (compare Fig 1E and F), and combined with Fig. 1B, seems to represent negligible changes, thus further supporting the stability of CME and its suitability for longitudinal studies.

### Using CMEs in a longitudinal aging experiment

The stability of CME over a prolonged period of time makes it a useful resource for longitudinal studies requiring a microbially-diverse yet stable source for host colonization. To test its usefulness, we used CMEs in an experiment aimed at characterizing the gut microbiome of aging worms. To this end, we raised worms on plates seeded with CME aliquots, transferring aging worms every two days away from their progeny to fresh plates with aliquots of the same stored CME, and sampling both lawns and worms for microbiome analysis.

Two independent experiments were performed, each with a separately prepared CME generated with distinct fruit enrichments. The resulting microbial composition of the two CMEs was accordingly very different (Fig. 2A and B). One was dominated by members of the families *Acetobacteraceae, Alcaligenaceae, Brevibacteriaceae, Moraxellaceae* and *Microbacteriaceae* (CME-1, Fig. 2A); the other, by *Alteromonadaceae, Bacillaceae, Crocinitomicaceae*, and *Verrucomicrobiaceae* (CME-2, Fig. 2A).

**Figure 2.**
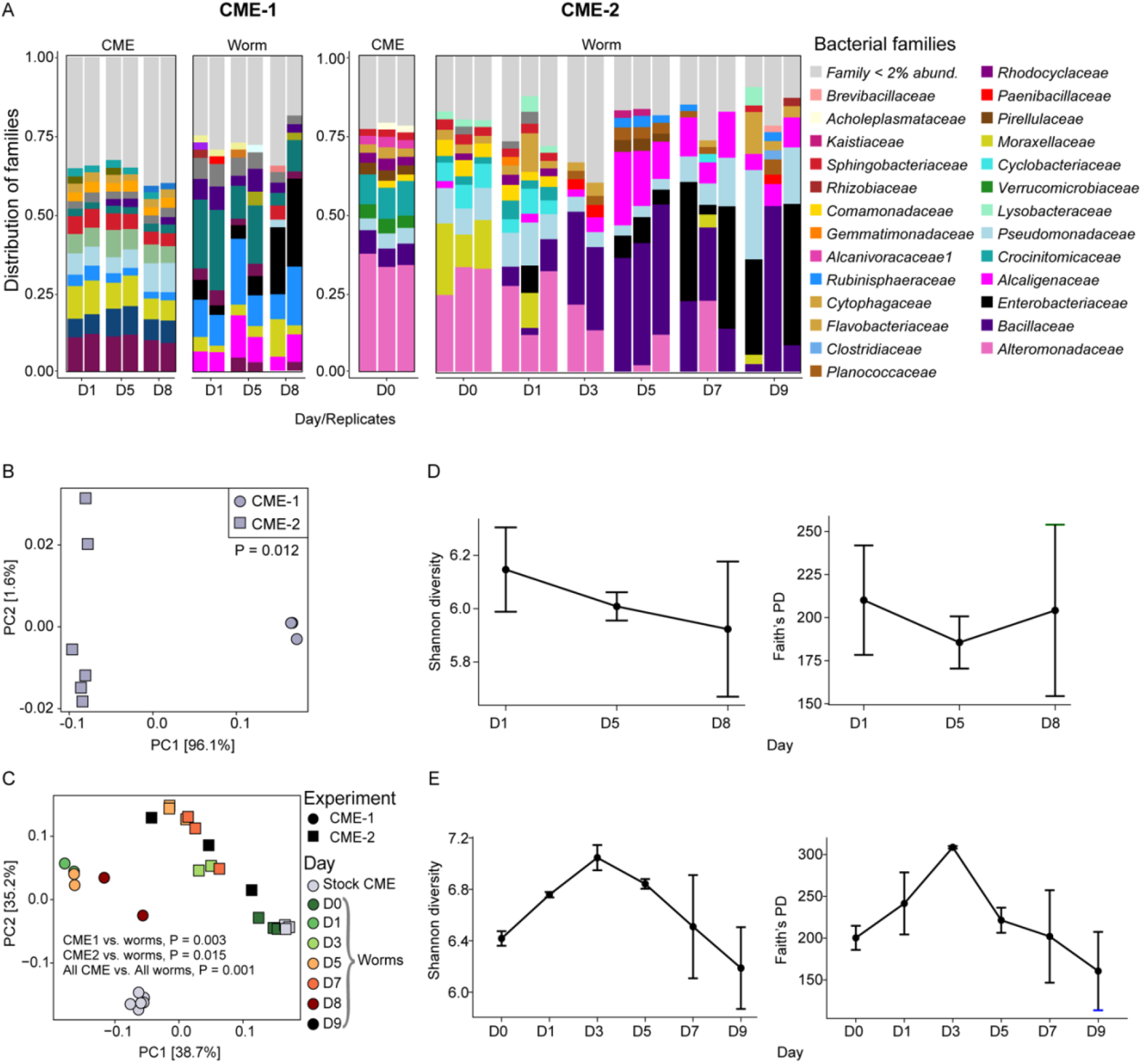
CME microbiome stability enables long-term aging experiments in C. elegans. **(A)** Relative abundance of bacterial families in CME stocks and in worm gut microbiomes. Each stacked bar represents an independent population of 50-100 worms and is color-coded by bacterial family. Shown for CME2 stock community is only one time point out of 6 with similar microbial composition as in CME1 (others are not shown). (**B**) Principal component analysis of CME1 and CME2 stock samples highlighting differences between them based on weighted UniFrac distances. P-value obtained with PERMANOVA (**C**) Principal component analysis based on weighted UniFrac distances of worm gut microbiomes during aging, sampled in designated time points and the respective CMEs at Day 0. (**D**) Alpha diversity indices for worm microbiomes raised on CME1 and sampled at designated days. Shown are means ± SD from independent samples (N = 2 plates per time point). (**E**) Alpha diversity indices for worms raised on CME2 (N = 3 per time point).

Previous work has shown that worms harbored a characteristic gut microbiome, distinct from their microbial environments but similar among worms raised in different environments [10,21,24]. This trend held true here as well (Fig. 2C). In addition, microbiome composition across time points revealed age-associated shifts in the worm gut community (Fig. 2C). Those differed between the two experiments but shared an age-dependent increase in the abundance of bacteria of the *Enterobacteriaceae* family (Fig. 2A), as observed previously [24]. Moreover, the gut bacterial diversity declined overall in aging worms (Fig. 2D and E), consistent with other studies reporting age-associated reduction in gut microbiome diversity [24,41].

## Discussion

Experiments using monocultures and synthetic communities have been invaluable for dissecting mechanisms of animal-microbe interactions. The CME-based approach described here complements these reductionist systems by enabling investigation of emergent microbial community properties within the host over time, including succession, resilience, and competitive dynamics.

In natural environments, microbial communities are highly dynamic, making it difficult to disentangle host-driven changes from environmental variation. By maintaining compositional stability during storage while preserving ecological complexity, CME reduces environmental variability and provides consistent microbial exposure across experiments. Together, these features establish a practical framework for longitudinal studies of host–microbiome interactions under controlled yet ecologically relevant conditions.

Our microbiome analysis in aging worms demonstrated age-related declines in microbial diversity and temporal changes in composition consistent with previous work [24], including a recurring increase in *Enterobacteriaceae* in older worms. The reproducibility of this pattern across CME-based experiments reinforces its robustness as an aging-associated signature. Notably, the consistent emergence of *Enterobacteriaceae* across monocultures, synthetic communities containing this family, and now naturally complex CME communities suggests that this shift is not an artifact of experimental simplification but a general feature of the aging *C. elegans* gut environment.

While the results described here use CMEs to focus on the host-associated microbiome, two additional studies utilized CMEs to study eco-evolutionary aspects of the worm gut microbiomes. In these studies, CMEs were useful to demonstrate that regardless of microbial diversity, stress-resistant *C. elegans* dauer larvae were largely devoid of gut bacteria, and that worm population growth can shape their respective microbial environments [6,34]. Together with the present study, these studies highlight the broad applicability of CME in microbiome research in relevant ecological contexts.

Although our experiments focused on *C. elegans*, CMEs could be used for other useful nematode models such as *P pacificus* [42] or adapted across more distant animal models. CMEs could be applied to modify environments or diets of other animals such as *Drosophila*, or in earthworm’s habitats, where they could contribute to studies of processes such as nutrient cycling or host-microbiome interactions. CMEs could also serve as natural microbiome “inoculum” for aquatic systems or animal-housing environments, to expose animal models to ecologically relevant microbial communities. Overall, CMEs provide a flexible and broadly applicable framework for studying host-associated microbiomes across biological systems of varying complexity.

## Acknowledgments

Work described in this manuscript was supported by NIH grants R01AG061302 and R01ES034012.

## Data Availability

Raw data is available in the NCBI SRA database with accession number PRJNA1462967.

### Code availability

R-scripts used to generate figures are available at https://rpubs.com/MicroRB/CME_Method_Paper

### Authors contributions

RB and MS conceived the project and wrote the manuscript. RB and RC performed the experiments and analyzing results; RB was further responsible for analyzing sequencing data.

## References

1. Levin D, Raab N, Pinto Y et al. Diversity and functional landscapes in the microbiota of animals in the wild. Science 2021;372(6539). 10.1126/science.abb5352.

2. Kolodny O, Callahan BJ, Douglas AE. The role of the microbiome in host evolution. Philos Trans R Soc Lond B Biol Sci 2020;375(1808). 10.1098/rstb.2019.0588.

3. Week B, Russell SL, Schulenburg H et al. Applying evolutionary theory to understand host–microbiome evolution. Nat Ecol Evol 2025;9(10):1769–1769. 10.1038/s41559-025-02846-w.

4. Shreiner AB, Kao JY, Young VB. The gut microbiome in health and in disease. Curr Opin Gastroenterol2015;31(1):69– 75. 10.1097/MOG.0000000000000139.

5. Johnke J, Zimmermann J, Stegemann T et al. Caenorhabditis nematodes influence microbiome and metabolome characteristics of their natural apple substrates over time. mSystems 2025;10(2). 10.1128/msystems.01533-24.

6. Bodkhe R, Sankaran K, Shapira M. Caenorhabditis elegans populations shape their microbial environment. npj Biofilms Microbiomes 2026; published online 4 Apr 2026. 10.1038/s41522-026-00975-z.

7. Ludington WB, Ja WW. Drosophila as a model for the gut microbiome. PLoS Pathog 2020;16(4):e e1008398. 10.1371/journal.ppat.1008398.

8. Barron AJ, Agrawal S, Lesperance DNA et al. Microbiome-derived acidity protects against microbial invasion in Drosophila. Cell Rep 2024;43(4):114087. 10.1016/j.celrep.2024.114087.

9. Sree Kumar H, Wisner AS, Refsnider JM et al. Small fish, big discoveries: zebrafish shed light on microbial biomarkers for neuro-immune-cardiovascular health. Front Physiol 2023;14:1186645. 10.3389/fphys.2023.1186645.

10. Zhang F, Berg M, Dierking K et al. Caenorhabditis elegans as a model for microbiome research. Front Microbiol2017;8:485. 10.3389/fmicb.2017.00485.

11. Douglas AE. Simple animal models for microbiome research. Nat Rev Microbiol 2019;17(12):764–764. 10.1038/s41579-019-0242-1.

12. Montalvo-Katz S, Huang H, Appel MD et al. Association with soil bacteria enhances p38-dependent infection resistance in Caenorhabditis elegans. Infect Immun 2013;81(2):514. 10.1128/IAI.00653-12.

13. Haçariz O, Viau C, Karimian F et al. The symbiotic relationship between Caenorhabditis elegans and members of its microbiome contributes to worm fitness and lifespan extension. BMC Genomics 2021;22(1):364. 10.1186/s12864-021-07695-y.

14. Pérez-Carrascal OM, Choi R, Massot M et al. Host preference of beneficial commensals in a microbially diverse environment. Front Cell Infect Microbiol 2022;12:795343. 10.3389/fcimb.2022.795343.

15. O’Donnell MP, Fox BW, Chao PH et al. A neurotransmitter produced by gut bacteria modulates host sensory behaviour. Nature 2020;583(7816):415. 10.1038/s41586-020-2395-5.

16. Manterola M, Palominos MF, Calixto A. The heritability of behaviours associated with the host gut microbiota. Front Immunol 2021;12:658551. 10.3389/fimmu.2021.658551.

17. Kiontke K, Sudhaus W. Ecology of Caenorhabditis species. WormBook 2006:1–14. 10.1895/wormbook.1.37.1.

18. Trang K, Bodkhe R, Shapira M et al. Compost microcosms as microbially diverse, natural-like environments for microbiome research in Caenorhabditis elegans. J Vis Exp 2022;(187):64393. 10.3791/64393.

19. Schulenburg H, Félix MA. The natural biotic environment of Caenorhabditis elegans. Genetics 2017;206(1):55–55. 10.1534/genetics.116.195511.

20. Frézal L, Félix MA. C. elegans outside the Petridish. eLife 2015;4:e05849. 10.7554/eLife.05849.

21. Berg M, Stenuit B, Ho J et al. Assembly of the Caenorhabditis elegans gut microbiota from diverse soil microbial environments. ISME J 2016;10(8):1998–1998. 10.1038/ismej.2015.253.

22. Dirksen P, Assié A, Zimmermann J et al. CeMbio—the Caenorhabditis elegans microbiome resource. G3 (Bethesda) 2020;10(9):3025. 10.1534/g3.120.401309.

23. Dirksen P, Marsh SA, Braker I et al. The native microbiome of the nematode Caenorhabditis elegans: gateway to a new host–microbiome model. BMC Biol 2016;14:25. 10.1186/s12915-016-0258-1.

24. Choi R, Bodkhe R, Pees B et al. An Enterobacteriaceae bloom in ageing animals is restrained by the gut microbiome. Aging Biol 2024;1(1):20240024. 10.59368/agingbio.20240024.

25. Zhang F, Weckhorst JL, Assié A et al. Natural genetic variation drives microbiome selection in the Caenorhabditis elegans gut. Curr Biol 2021;31(12):2603–2603.e9. 10.1016/j.cub.2021.04.046.

26. Koyle ML, Veloz M, Judd AM et al. Rearing the fruit fly Drosophila melanogaster under axenic and gnotobiotic conditions. J Vis Exp 2016;(113):54219. 10.3791/54219.

27. Correa MA, Matusovsky B, Brackney DE et al. Generation of axenic Aedes aegypti demonstrates live bacteria are not required for mosquito development. Nat Commun 2018;9:2477. 10.1038/s41467-018-07014-2.

28. Kešnerová L, Mars RAT, Ellegaard KM et al. Disentangling metabolic functions of bacteria in the honey bee gut. PLoS Biol 2017;15(12):e2003467. 10.1371/journal.pbio.2003467.

29. Jennings SA V, Clavel T. Synthetic communities of gut microbes for basic research and translational approaches in animal health and nutrition. Annu Rev Anim Biosci 2026. 10.1146/annurev-animal-021022.

30. Gallet A, Halary S, Duval C et al. Disruption of fish gut microbiota composition and holobiont metabolome during a simulated Microcystis aeruginosa bloom. Microbiome 2023;11:185. 10.1186/s40168-023-01558-2.

31. Jackrel SL, Broe TY. Intraspecific variation in leaf litter alters fitness metrics and the gut microbiome of consumers. Oecologia 2023;202(4):769–769. 10.1007/s00442-023-05435-5.

32. Sun C, Wang Y, Jia C et al. Which is more priority, substrate type or food quality? A case study on a tropical coral reef sea cucumber Stichopus chloronotus. BMC Microbiol 2024;24:142. 10.1186/s12866-024-03670-1.

33. Muslim A, Mizushima D, Phumee A et al. Tannic acid shaped microbiome composition in midguts and rearing microcosms of Aedes triseriatus. Front Microbiol 2026;17:1755894. 10.3389/fmicb.2026.1755894.

34. Bodkhe R, Trang K, Hammond S et al. Emergence of dauer larvae in Caenorhabditis elegans disrupts continuity of host–microbiome interactions. FEMS Microbiol Ecol 2024;100(12):fiae149. 10.1093/femsec/fiae149.

35. Callahan BJ, McMurdie PJ, Rosen MJ et al. DADA2: high-resolution sample inference from Illumina amplicon data. Nat Methods 2016;13(7):581– =3. 10.1038/nmeth.3869.

36. Quast C, Pruesse E, Yilmaz P et al. The SILVA ribosomal RNA gene database project: improved data processing and web-based tools. Nucleic Acids Res 2013;41(Database issue). 10.1093/nar/gks1219.

37. McMurdie PJ, Holmes S. phyloseq: an R package for reproducible interactive analysis and graphics of microbiome census data. PLoS One 2013;8(4):e61217. 10.1371/journal.pone.0061217.

38. Wright ES. Fast and flexible search for homologous biological sequences with DECIPHER v3. R J 2024;16(2):191–191. 10.32614/rj-2024-022.

39. Schliep KP. phangorn: phylogenetic analysis in R. Bioinformatics 2011;27(4):592. 10.1093/bioinformatics/btq706.

40. Wickham H. ggplot2: Elegant Graphics for Data Analysis. Cham: Springer, 2016. 10.1007/978-3-319-24277-4.

41. Deng F, Li Y, Zhao J. The gut microbiome of healthy long-living people. Aging (Albany NY) 2019;11(2):289. 10.18632/aging.101771.

42. Lo WS, Sommer RJ, Han Z. Microbiota succession influences nematode physiology in a beetle microcosm ecosystem. Nat Commun 2024;15:49513. 10.1038/s41467-024-49513-5.

